# The Rise of Open Data Practices Among Bioscientists at the University of Edinburgh

**DOI:** 10.1101/2024.02.18.580901

**Authors:** Haya Deeb, Suzanna Creasey, Diego Lucini de Ugarte, George Strevens, Trisha Usman, Hwee Yun Wong, Megan A. M. Kutzer, Emma Wilson, Tomasz Zieliński, Andrew J. Millar

## Abstract

Open science promotes the accessibility of scientific research and data, emphasising transparency, reproducibility, and collaboration. This study assesses the openness and FAIRness (Findable, Accessible, Interoperable, and Reusable) of data-sharing practices within the biosciences at the University of Edinburgh from 2014 to 2023. We analysed 555 research papers across biotechnology, regenerative medicine, infectious diseases, and non-communicable diseases. Our scoring system evaluated data completeness, reusability, accessibility, and licensing, finding a progressive shift towards better data-sharing practices. The fraction of publications that share all relevant data increased significantly, from 7% in 2014 to 45% in 2023. Data involving genomic sequences were shared more frequently than image data or data on human subjects or samples. The presence of data availability statement (DAS) or preprint sharing correlated with more and better data sharing, particularly in terms of completeness. We discuss local and systemic factors underlying the current and future Open data sharing. Evaluating the automated ODDPub (Open Data Detection in Publications) tool on this manually-scored dataset demonstrated high specificity in identifying cases where no data was shared. ODDPub sensitivity improved with better documentation in the DAS. This positive trend highlights improvements in data-sharing, advocating for continued advances and addressing challenges with data types and documentation.

## Introduction

Open science is the movement aimed at making scientific research, data, and dissemination more universally accessible. It encompasses a range of practices that promote transparency, reproducibility and collaboration, including open access to publication, open peer review, and open data (1). Open data, in particular, is a critical component of open science, as it ensures that the data underlying research findings are freely available for scrutiny, validation, and reuse by others in the scientific community (2). By making data openly available, researchers can enhance the reproducibility of their findings, facilitate discoveries through data reuse, and promote greater transparency in scientific research (3,4).

The concept of FAIR data – emphasising data that is ‘Findable’, ‘Accessible’, ‘Interoperable’, and ‘Reusable’ –(5,6) was introduced to promote the transparency and integrity of shared research data, even if those data are not fully Open. While the availability of open data is a positive step towards transparency, the utility of this data often depends on how well it adheres to the FAIR principles (6). Simply being ‘open’ does not guarantee that data are well-organised, thoroughly documented, or easily usable. Furthermore, while access to certain datasets may be restricted due to privacy or confidentiality concerns, these datasets can still adhere to FAIR principles. This means that even though the data is not openly accessible, it could still be findable, accessible under certain conditions, interoperable, and reusable within those constraints (7).

In recent years, there has been a significant shift in policies regarding data sharing. Major funding agencies and journals have increasingly mandated that researchers share the data underlying their publications. For instance, the National Institutes of Health (NIH) and the European Commission have implemented policies that require data sharing as a condition of funding (8,9). Journals such as PLOS ONE and Nature also require authors to make their data publicly available upon publication (10,11). These policy changes reflect a growing recognition of the value of open data in advancing scientific knowledge.

Despite the benefits and policy support, several challenges persist in data sharing. Researchers may be reluctant to share data due to concerns about data misuse, loss of competitive advantage, or the significant time and effort required to prepare data for sharing (12). Additionally, issues such as data privacy, and the lack of standardised formats and repositories can complicate data-sharing efforts (13–15). Intellectual property rights and considerations surrounding human data disclosure pose consistent challenges (16,17). Though data anonymisation has been advanced as a solution, risks of re-identification remain, especially among vulnerable populations (14,15). Additionally, the public sharing of anonymised data may not always be covered by the original consent agreements, adding a layer of complexity to privacy protection.

Bioscience research exemplifies the balancing act between open access and data protection. the University of Edinburgh updated its Research Data Management in 2021 (18). It had evolved to include adherence to FAIR principles and GDPR compliance, highlighting the importance of data accessibility, interoperability, privacy, and security. This progression mirrors the broader movement towards more transparent, accessible, and collaborative research practices. Therefore, our research aims to assess the openness and FAIRness (Findability, Accessibility, Interoperability, and Reusability) of data-sharing practices within the biosciences at the University of Edinburgh from 2014 to 2023. We seek to provide a detailed analysis of how different research areas— biotechnology, regenerative medicine, infectious diseases, and non-communicable diseases— perform in terms of data sharing. Additionally, we investigate the correlation between Data Availability Statements (DAS) and preprint status on the quality of data sharing. By highlighting the current state of data-sharing practices and identifying areas for improvement, this study aims to contribute to the ongoing efforts to promote transparency and reproducibility in scientific research.

## Methods

### Definition of Data

The term “Data” refers to both qualitative and quantitative information necessary to reproduce research results. This includes primary data, such as raw experimental results; secondary data, like derived analytical values, and the software code used to derive those values. Data also includes images and videos used for conducting analyses. When quantitative analysis is performed on images, these images are considered raw data, and the resulting numerical values are considered processed data. Images and videos used solely for illustrative purposes are excluded.

### Sampling Framework and Selection Process

Research groups within the biosciences at the University of Edinburgh were selected for study, based on the educational interests of five student researchers. 57 groups were selected from the University’s public research websites in the following areas: Non-Communicable Diseases (NCDs), Infectious Diseases and their treatments (InfD), Microbial Biotechnology (BioTech), Stem Cells and Regeneration (SRM). In each area, all research groups in the School of Biological Sciences were selected. Research groups in the same areas based in the College of Medicine were included, forming 17 out 57 research groups. This allowed each student to score and analyse around 100 articles within their project time limit.

Articles from each group were retrieved from the University’s public Edinburgh Research Explorer website (https://www.research.ed.ac.uk/) (19), which is a curated, institutional research information system. To be included a group needed a substantial publication record, with at least 10 journal articles published between 2014 and 2023. From each selected group, we randomly chose journal articles for each year within our study period to ensure a representative sample. Our focus was strictly on original research papers that generated new datasets, excluding other publication types such as reviews, editorials, and commentaries. This selection process was carried out by five biology students, who manually reviewed and verified each article based on shared scoring criteria, that were documented on an internal wiki site. The selected articles and scores were compiled on spreadsheets and combined, with each article categorised by research areas NCDs, InfD, Biotech, and SRM.

### Scoring Criteria

The scoring system was developed based on those established by Roche et al. (14) and the FAIR Data Self-assessment tool by the Australian Research Data Commons (20). We tailored the scoring criteria, pertinent to our study’s objectives. A key adaptation involved refining the scoring scale from five levels, as used by Roche et al., to four. This reduction was motivated by a need to distinctly address file format and metadata quality without conflating them with raw versus processed data considerations, thereby eliminating potential overlap and ambiguities in evaluations.

While Roche et al. (14) sampled from a popular data repository Dryad, where sampling started from archived datasets, this study involved selecting articles while being blinded as to whether any data was shared. When articles reused previously published data, a similar approach to Hamilton et al. (21) was taken. The four criteria – ‘Completeness’, ‘Reusability’, ‘Accessibility’, and ‘Licence’ – were established to assess independent aspects of data sharing. The Completeness score was based upon the whole article. The criteria for the quality of sharing, Reusability, Accessibility and Licence, were assessed only for the best-shared dataset, in contrast to Roche et al. (14), to avoid confounding these quality criteria with the Completeness of sharing.

1. **Data Completeness:** To evaluate completeness, we assigned scores from 1 to 4, with 4 indicating complete data sharing necessary for reproducing all analyses and results. The highest score did not require sharing the partially processed data at each analysis step, provided that the raw data and sufficient description of data processing steps were available. Scores of 2 and 3 indicated minor and major data omissions, respectively. This ranking accommodated the wide variety of data types and volumes in individual articles and between articles in our sample (see Discussion), as in Roche et al. (14). When a score of 1 was assigned, indicating no data sharing, the subsequent criteria were also assigned a score of 1 (Not Scored). The absence of shared data negated the possibility of evaluating reusability, accessibility, and licence clarity. We assessed the completeness of data sharing, irrespective of its location - whether in texts, figures, tables, supplementary materials, or external repositories (Table 1).
2. **Data Reusability:** Reusability scores, ranging from 1 to 4, were based on data being in a non-proprietary, human- and machine-readable format, and accompanied by informative metadata. A score of 4 required fulfillment of all these aspects. The criteria for classifying files according to their non-proprietary, human- or machine-readable format are detailed in Table S1 of the supplementary material file 1, adapted from Roche et al (14). The inclusion of metadata in this criterion highlights its indispensable role in preserving the data’s long-term utility (Table 1). Proprietary formats were penalised unless they were community-accepted or mandated by public repositories (24,25).
3. **Data Accessibility:** Accessibility was assessed on whether the data was accessible, had a unique or persistent identifier (PID) and was archived in a public repository. Scores ranged from 1 to 4, with 4 indicating fulfillment of all conditions. Data with a PID in a public repository but inaccessible or requiring permission received the lowest score, reflecting our focus on data openness. Scoring details are in Table 1.
4. **Data Licence:** The licence type was scored from 1 to 4. Open licences received the highest score, restrictive licences scored 3, and data without a specified licence or governed by the article’s licence scored 2. Criteria are detailed in Table 1.

**Table 1:** Scoring Criteria used for the Assessment.

| Data Completeness |  |  |
| --- | --- | --- |
| Score | Description | Criteria |
| 4 | Exemplary | All the data necessary to reproduce the analyses and results (in practice) are present within the article (e.g. as supplementary information, figures) or archived in external repositories. Both raw and processed datasets from all methods utilised and mentioned in the study are provided. |
| 3 | Good | Most of the data necessary to reproduce the analyses and results (in practice) are present within the article (e.g. as supplementary information, figures) or are archived in external repositories.<br>Processed data from all methods utilised and mentioned in the article are provided, lacking only a small amount of raw datasets. |
| 2 | Large Omission | Main analyses in the paper cannot be redone because essential datasets are missing AND/OR only summary statistics (e.g. means, standard deviation) obtained from methods utilised and mentioned in the article are archived, no raw data provided. |
| 1 | Poor | Neither processed nor raw data are archived/present in the article<br>OR the incorrect data are archived. |

| Reusability |  |  |
| --- | --- | --- |
| Score | Description | Criteria (For best dataset) |
| 4 | Exemplary | Good formats and metadata. Data is archived in a (1) non-proprietary, (2) human- and machine-readable file format that facilitates data aggregation and can be processed with both free and proprietary software (e.g. csv, text); other formats were allowed as *de facto* community standards (see Table S1, Supplementary Material File 1). (3) Highly informative metadata (such that column headings, abbreviations, and units can be understood in isolation from the original paper). |
| 3 | Good | Data is archived in (1) a non-proprietary OR (2) a human- and machine-readable file format that facilitates data aggregation and can be processed with both free and proprietary software. OR (3) Metadata must at least be sufficiently informative to be understood when combined with the paper. |
| 2 | Average | Poor formats and metadata. Data is archived in (1) a proprietary OR (2) human- but not machine-readable file format (e.g., pdf, jpeg). AND (3) Metadata is not sufficiently informative when combined with the paper. |
| 1 | Poor | Not Scored |

| Accessibility |
| --- |

| Score | Description | Criteria (For best dataset) |
| --- | --- | --- |
| 4 | Exemplary | Data is accessible, has a Persistent Identifier (PID) assigned or a unique identifier from a high-tier repository (such as a GenBank genInfo number), and is stored in an online public repository (e.g. Figshare). |
| 3 | Good | Data is accessible, has either a PID or a unique identifier from a high-tier repository OR is stored in an online public repository. |
| 2 | Average | Data is accessible but does not have a PID or unique identifier and is not stored in an online public repository (e.g. data shared in the supplementary information section of the article). |
| 1 | Poor | Not Scored |

| Licence |  |  |
| --- | --- | --- |
| Score | Description | Criteria (For best dataset) |
| 4 | Exemplary | Data has a permissive licence (e.g. CC0, CC-BY) and code has an Open Software licence (if applicable). |
| 3 | Good | Licence(s) are present but not all are permissive. Data has a restrictive licence (e.g. CC-BY-SA/NC) and/or code has a Closed Software licence (if applicable). |
| 2 | Average | No explicit licence is provided for data or software. |
| 1 | Poor | Not Scored |

### Additional Data Evaluation Criteria

Our study assessed the sharing of different data types in biosciences research, using a consistent scoring system. For each data type—image, genomic, and human—scoring was as follows: ‘Not Applicable’ (NA) if the data type was irrelevant to the study, ‘Not Shared’ (0) if it was relevant but not shared, and ‘Shared’ (1) if it was both relevant and fully archived. This system applied to unprocessed images for potential re-analysis, nucleotide sequence-based assays including transcriptomics for genomic data, and data involving human stem cell lines or human participants for human data. We further evaluated the storage of shared data with a separate variable, ‘Storage’, which identified the location of data sharing: ‘Repository’ if data was shared in an external repository, ‘Supplementary Material’ if in the publication’s supplementary materials, ‘Both’ for data shared in both locations, ‘DOI in Methods’ for cases where only the dataset DOI is cited in the methods section without providing the project’s actual dataset, and ‘NA’ for instances where no data was shared. Additionally, we tracked the publication of corresponding preprints, utilising Google Scholar, medRxiv and bioRxiv preprint servers for verification.

Data Availability Statements (DAS) in each article were only recognised if explicitly titled as such or with similar designations. DAS were categorised as ‘Shared’ when indicating unrestricted data access, or ‘Not Shared’ when access was conditional. Articles without a specific DAS section were marked as ‘Not Presented’. We assume that the inclusion of a DAS reflects journal policy more than researcher preference (20). Journal policies were not scored directly, because the historic policy versions that applied upon publication back to 2014 were not reliably available. Finally, we assessed the use of analytical software in the research by recording whether it was mentioned (’Yes’ or ‘No’). Additionally, for analyses that required coding, we evaluated whether the code was shared (’Shared’), used but not shared (’Used and Not Shared’), or not applicable (’NA’), based on the documentation within the publication.

### Data Extraction and Quality Control Assessment

The collected data, including both scoring assessments and supplementary variables, were cataloged in an Excel spreadsheet. To test the reliability of our assessment, 15% of the papers from each evaluator’s dataset, from a random selection supplemented to cover the entire spectrum of scoring values and years, was re-scored by another researcher. This blind reassessment returned a congruence in scoring assessments between the original and secondary evaluations from 86% to 90%. Consequently, the initial assessments were retained for final analysis.

### Integration of ODDPub Algorithm for Detection the Open Data Practices

We applied the Open Data Detection in Publications (ODDPub) text mining algorithm to further evaluate the findings and to explore the automation of open science practice assessment across our dataset. Originally developed for and applied in biomedical datasets, as documented in Riedel et al. (22), ODDPub assists in systematically identifying open data and code practices from full text documents of research articles. We retrieved publication full texts in PDF format from DOIs using the Find Full Text function in EndNote 20 (RRID:SCR_014001) or hand searching. We converted PDF files to text format using the pdftools R package (23) for processing by ODDPub, via the ODDPub R Package. ODDPub provided several key outputs: *is_open_data* is a binary indicator noting whether open data was detected, and *open_data_category* indicates the type of data sharing (e.g. supplement, general-purpose repository, field-specific repository), or records ‘NA’ if no data is found. Additionally, *is_open_code* denotes whether open code was detected (TRUE/FALSE).

Supplementary to these, *open_data_statements* and *open_code_statements* extract text related to their respective sharing details, recording ‘NA’ if no reference is made to sharing in the text.

To evaluate the performance of ODDPub in our context, we calculated its sensitivity and specificity. Sensitivity, or true positive rate, measures the proportion of actual positives correctly identified by the algorithm, while specificity, or true negative rate, measures the proportion of negatives correctly identified (Supplementary material file 2). These metrics were used to compare ODDPub’s automated detection results with our manual assessment.

### Statistical Analysis

The descriptive statistics were presented as frequency and percentage. An ordinal regression model was used to test for changes over time in the four scoring criteria from 2014 to 2023. Additionally, ordinal regression analysis measured the influence of research area and other sharing variables such DAS, and preprint status on the scoring criteria. To account for potential intra-group similarities in data sharing practices, random effects for research groups were included in the regression models, positing that research groups may exhibit consistent data sharing behaviors. Significance was set at p < 0.05, and results were expressed as odds ratios with corresponding 95% confidence intervals. Additionally, thorough checks were conducted to validate the assumptions underlying each statistical model used in this analysis. Details of these assumption checks, along with the complete analytical methodology, have been documented and are available on Zenodo (https://doi.org/10.5281/zenodo.14169649). The analyses were performed using R and RStudio Software (Version 4.2.2).

## Results

### Paper Characteristics

From 2014 to 2023, the School of Biological Sciences at the University of Edinburgh published 3706 research papers, as recorded in the Edinburgh Research Explorer (19). Our study analysed 555 papers, representing approximately 15% of the total publications. This sample was divided into four research areas, with each area comprising between 20% and 35% of the sample (Table 2).

**Table 2:** Characteristics of the publications in the sample.

| Variable |  | Bio-Technolog<br>y (BioTech) | Infectious<br>Diseases<br>(InfD) | Non-Communica<br>ble<br>diseases<br>(NCD) | Stem Cells<br>and<br>Regeneratio<br>n<br>(SRM) | Total |
| --- | --- | --- | --- | --- | --- | --- |
|  |  | (n= 130) | (n=198) | (n=121) | (n=106) | (n=555) |
| Preprint | Yes | 20 (15.38%) | 48 (24.24%) | 31 (25.62%) | 25 (23.58%) | 124<br>(22.34%) |
|  | No | 110<br>(84.62%) | 150 (75.76%) | 90 (74.38%) | 81 (76.42%) | 431<br>(77.66%) |
| DAS | Open<br>Data | 47 (36.15%) | 75 (37.88%) | 47 (38.84%) | 47 (44.34%) | 216<br>(38.92%) |
|  | Restrict<br>ed Data | 9 (6.92%) | 21 (10.60%) | 21 (17.36%) | 14 (13.21%) | 65 (11.71%) |
|  | Not<br>presente | 74 (56.92%) | 102 (51.52%) | 53 (43.80%) | 45 (42.45%) | 274<br>(49.37%) |
|  | d |  |  |  |  |  |
| Analysis Program | Yes | 97 (74.62%) | 183 (92.42%) | 120 (99.17%) | 92 (86.79%) | 492 (88.65%) |
|  | No | 33 (25.38%) | 15 (7.58%) | 1 (0.83%) | 14 (13.21%) | 63 (11.35%) |
| Code Sharing | Used and Shared | 17 (13.08%) | 24 (12.12%) | 37 (30.58%) | 34 (32.08%) | 112 (20.18%) |
|  | Used and not Shared | 3 (2.30%) | 29 (14.65%) | 19(15.70%) | 28 (26.42%) | 79 (14.23%) |
|  | Not Used | 110 (84.62%) | 145 (73.23%) | 65 (53.72%) | 44 (41.51%) | 364 (65.59%) |
| Data-Sharing | Not Shared (Completeness=1) | 6 (4.62%) | 23 (11.62%) | 26 (21.49%) | 24 (22.64%) | 79 (14.23%) |
|  | Data Shared (Completeness>1) | 124 (95.38%) | 175 (88.38%) | 95 (78.51%) | 82 (77.36%) | 476 (85.77%) |

We also observed varying levels of openness in research practices. Regarding the Data Availability Statement, approximately half of the papers did not present this statement in their publication. Most of the papers (492/555) shared the used analysis programs. Notably, among the 191 papers that required code for analysis, 112 actively shared their code, indicating a strong tendency towards openness in computational research. However, our findings show limited adoption of preprints, with only 124 out of 555 papers (22%) having associated preprints (Table 2).

### Assessment of Openness and FAIRness Criteria in Research Publications

The analysis of data-sharing practices among the four research areas reveals distinct trends. Overall, 86% of the publications shared some or all of their data (Completeness Score >1), with Biotechnology having the highest percentage of papers sharing data (95%). Infectious Diseases followed with 88% sharing data, while Non-Communicable Diseases and Regenerative Medicine had the lowest rates at 79% and 77% respectively (Table 1).

In the Completeness criterion (Figure 1), only 105/555 of the publications shared all available data, achieving a score of 4. Most articles (66%) received scores of 2 or 3, indicating moderate data sharing, while 14.2% of the publications scored 1, meaning no data were shared. The 79 articles with a Completeness score of 1 were not scored for the other criteria (where 1 indicates Not Scored).

**Figure 1:**
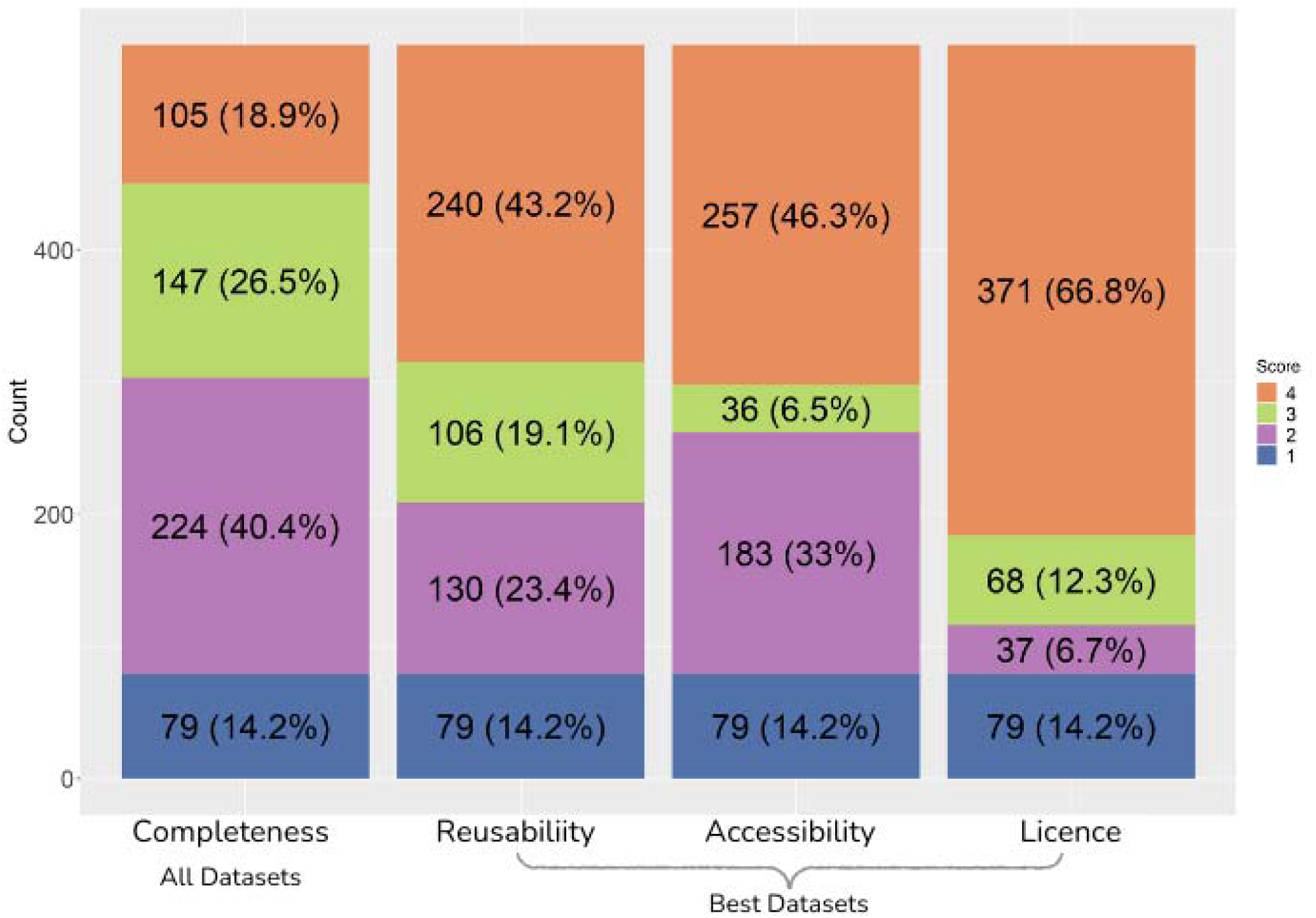
Scores of the four Criteria (Completeness, Reusability, Accessibility, and Licence) in all the research publications (n=555). Completeness is scored for all datasets required to reproduce the article, from 1 (no Open data) to 4 (all data are Open). The other criteria only score the best-shared dataset in each article, to avoid confounding these criteria with Completeness. For scoring details, please see Methods.

The quality of data sharing in the remaining 476 articles was assessed by scoring the best-shared data set, which was often shared very well. The largest segments of publications scored the maximum (4) for Reusability and Accessibility, with 43% and 46%, respectively. Permissive licensing was even more widespread, with fully 67% of publications having a permissive license for their shared data and (if applicable) an open software license (Figure 1).

### Decadal improvement in sharing, and regression models

Plotting the scores over the past ten years revealed a gradual improvement across all four criteria (Figure 2). The percentage of publications achieving a completeness score of 4 increased significantly, rising from 7% in 2014 to 45% in 2023. Similarly, the other criteria—Reusability,

**Figure 2:**
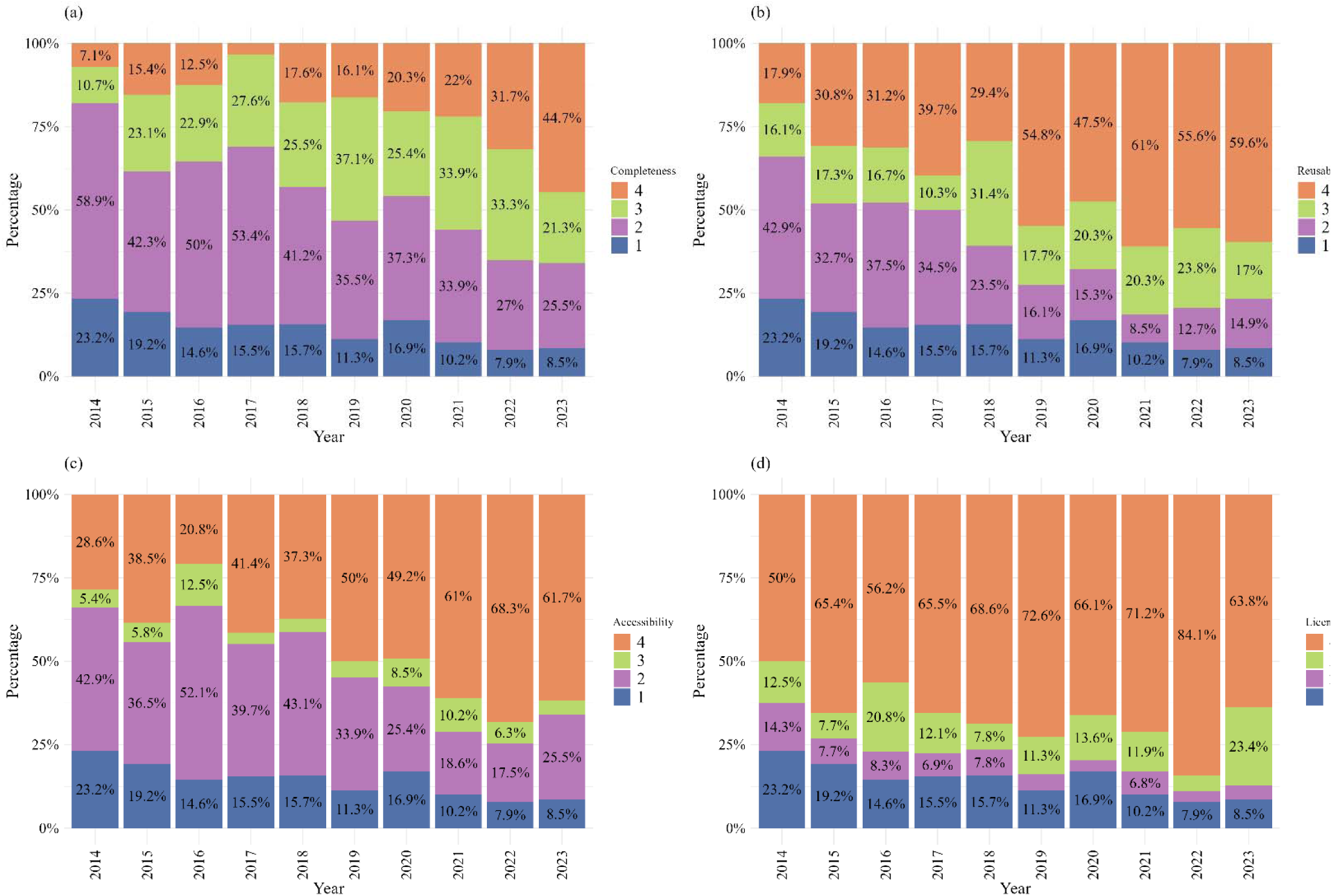
The distribution of scoring criteria over ten years. Increasing (a) Completeness, (b) Reusability, (c) Accessibility and (d) Licence are shown in the scores of n = 47-62 research publications per year.

Accessibility, and License—also showed notable improvements over the years, with publications achieving higher scores. These improvements were statistically significant according to the ordinal regression models (Table S2, Supplementary Material File 1) and remained significant when the research groups were included as a random effect in the regression model.

### Effect of the research area on the regression model

The distribution of scores varies among the four research areas. The articles in Non-Communicable Disease research show a higher frequency of lower completeness scores (Figure S1), and articles in Regenerative Medicine achieve more top scores in reusability. Ordinal regression analysis (Table S3, Supplementary Material File 1) confirms that Non-Communicable Disease articles show significantly lower completeness scores, when compared to Biotechnology as the reference research area, indicating substantial gaps in data sharing. Regenerative Medicine papers demonstrate higher reusability scores, suggesting better practices in data formatting and description. The Accessibility scores do not differ significantly across the research areas and Licence scores are high in all areas, so variation between them is not a pressing concern.

Our approach selects articles to study from the publication lists of research groups that focus on each research area, rather than assigning each article to a research area. Patterns that emerged at the research area level might therefore have reflected variations in data sharing among the groups, rather than varying practices across research areas. However, the findings above remain significant when the research groups are included as a random effect in the statistical models (Table S3, Supplementary Material File 1), consistent with differences in data sharing across research areas.

### Comparison of Data-Sharing Practices Across Research Areas

The comparison of data-sharing practices across four research areas reveals distinct patterns in the utilisation and sharing of genomic data. Among the papers that used genomic data, Regenerative Medicine stands out, with 94.3% actively sharing this data. In contrast, only 50% of the Biotechnology papers that utilised genomic data (27 out of 54) shared it. The research areas of Infectious Diseases and Non-Communicable Diseases demonstrated moderate sharing rates, with 74.2% (72 out of 97) and 58.3% (42 out of 72) of their genomic-using papers sharing the data, respectively (Table S4, Supplementary Material File 1).

For image data, Regenerative Medicine again leads with 29.2% of papers sharing images. Biotechnology, Infectious Diseases, and Non-Communicable Disease papers share image data at lower rates. Human data sharing is less common across all research areas, with Infectious Diseases leading at 14 out of 43 papers. Non-Communicable Disease and Regenerative Medicine papers have a lower rate of sharing human data, while the Biotechnology papers in our sample do not use human data (Table S4, Supplementary Material File 1).

In terms of data storage, the article’s supplementary materials were the most common storage location across all research areas, particularly in Biotechnology (53.1%) and Infectious Disease (37.9%). Regenerative Medicine showed a preference for using both repositories and supplementary materials, with 45.3% (Table S4, Supplementary Material File 1).

### Effectiveness and Accuracy of ODDPub in detecting Open Data Practice

The effectiveness of the ODDPub text mining algorithm in identifying open data and code sharing practices was evaluated by comparing its results against a manually assessed dataset. ODDPub employs a dictionary approach to identify terms relevant to data and code sharing and assigns publications binary TRUE/FALSE scores for several criteria (22). Figure 3 illustrates how ODDPub outcomes for open data and open code are distributed compared to manual Completeness and code sharing criteria.

**Figure 3:**
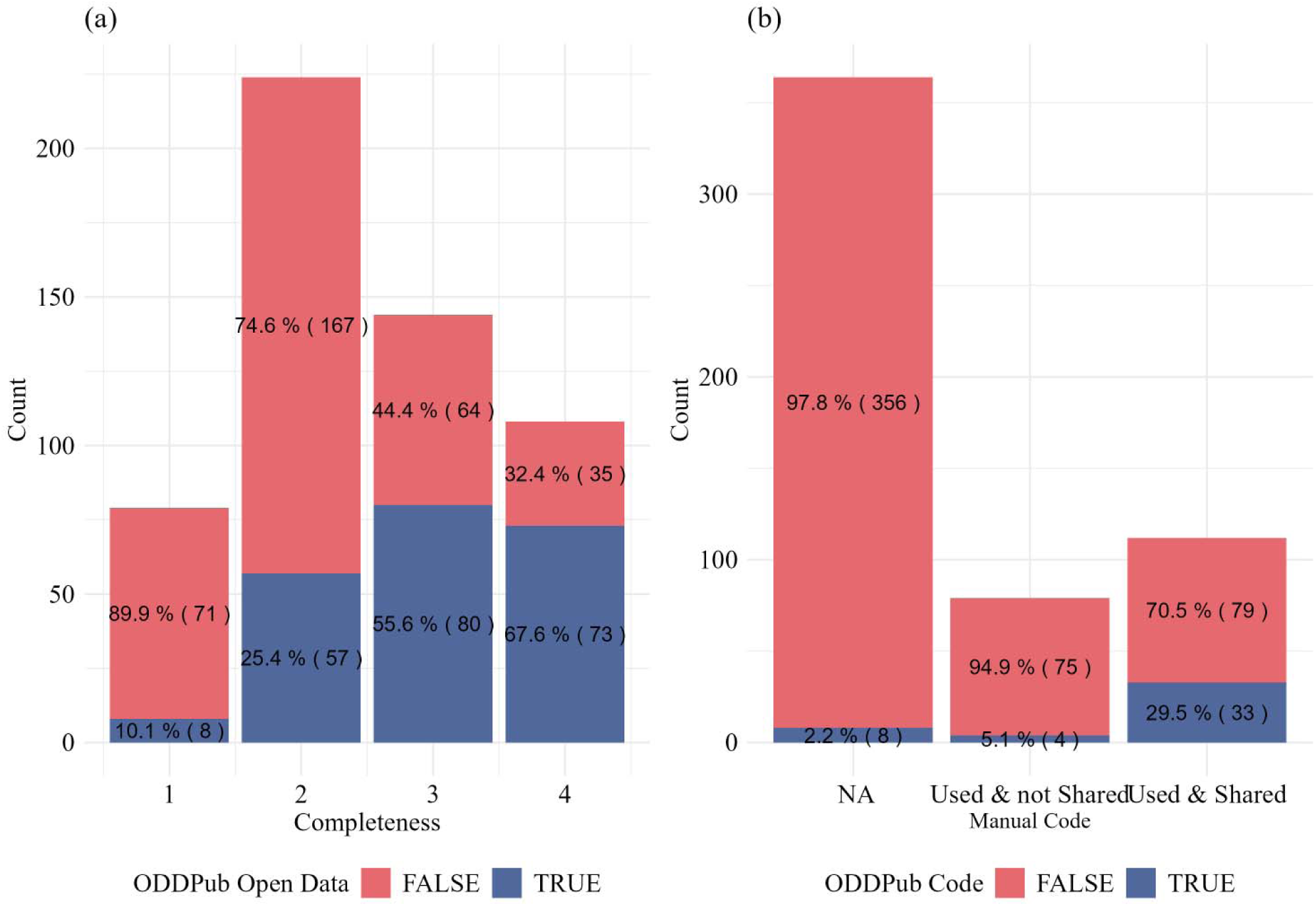
Manual and automated evaluation of Open Data and Open Code. (a) Distribution of ODDPub Open Data scores (Is_Open_Data, True/False) across manually-scored Completeness in all articles (N=555). (b) Comparison of Open Code evaluation between the Manual assessment and ODDPub Open Code score (Is_Open_Code, True/False).

Initially, ODDPub’s performance in detecting open data was assessed solely against our ‘completeness’ criterion, showing a sensitivity of 44% and a specificity of 90%. However, ODDPub scoring of Open data considers multiple factors, so we aligned the algorithm’s full definition of open data with our four manual criteria (detailed in Supplementary Material File 2). This improved the sensitivity to 52% while maintaining a high specificity of 90%. These metrics indicate the algorithm’s strong capability to identify non-open data instances. However, it shows limited effectiveness in identifying all the actual cases of open data.

In terms of open code practices, the algorithm recorded sensitivity and specificity rates of 30% and 97%, respectively. These results highlight ODDPub’s challenges in detecting true positive cases of open code while reaffirming its proficiency in ruling out instances where code is not shared.

### Impact of Data Availability Statement (DAS) and Preprint Status on Scoring Criteria

The ordinal regression analysis reveals significant associations between the presence of a Data Availability Statement (DAS) and preprint status on various data-sharing criteria (Table S5, Supplementary Material File 1). Papers with a ‘Shared’ DAS showed significantly higher scores across the four criteria. Papers with a DAS indicating data available upon request (‘Not Shared’) showed a smaller increase in Reusability, Accessibility, and License, but no significant change in Completeness. The presence of a preprint also significantly increased the odds of higher Completeness scores (Table S5, Supplementary Material File 1), though its effect on Reusability, Accessibility, and License clarity was not statistically significant. These findings underscore the positive correlations among data-sharing practices.

### DAS as an indicator of data sharing and its impact on ODDPub detection accuracy

Figure 4 illustrates the same correlation with Data Availability Statements (DAS), from the converse analysis, stratified by the Completeness score. Papers with higher completeness scores often had a ‘Shared’ DAS. Specifically, 88 papers with a completeness score of 4 had a ‘Shared’ DAS, while only 17 papers with a score of 4 had no DAS presented. As completeness scores decreased, the likelihood of lacking a DAS increased. However, 21 articles that indicated data were available upon request (a ‘Not Shared’ DAS) had in fact shared most of their data openly (Completeness 3), and articles with a ‘Shared’ DAS were represented in all Completeness categories. At the extreme, ten papers that claimed all data were available in their DAS paradoxically received a Completeness score of 1, indicating that no data were shared. These papers were published between 2014 to 2021. This discrepancy highlights a gap between the declared availability of data and the actual sharing of complete datasets.

**Figure 4:**
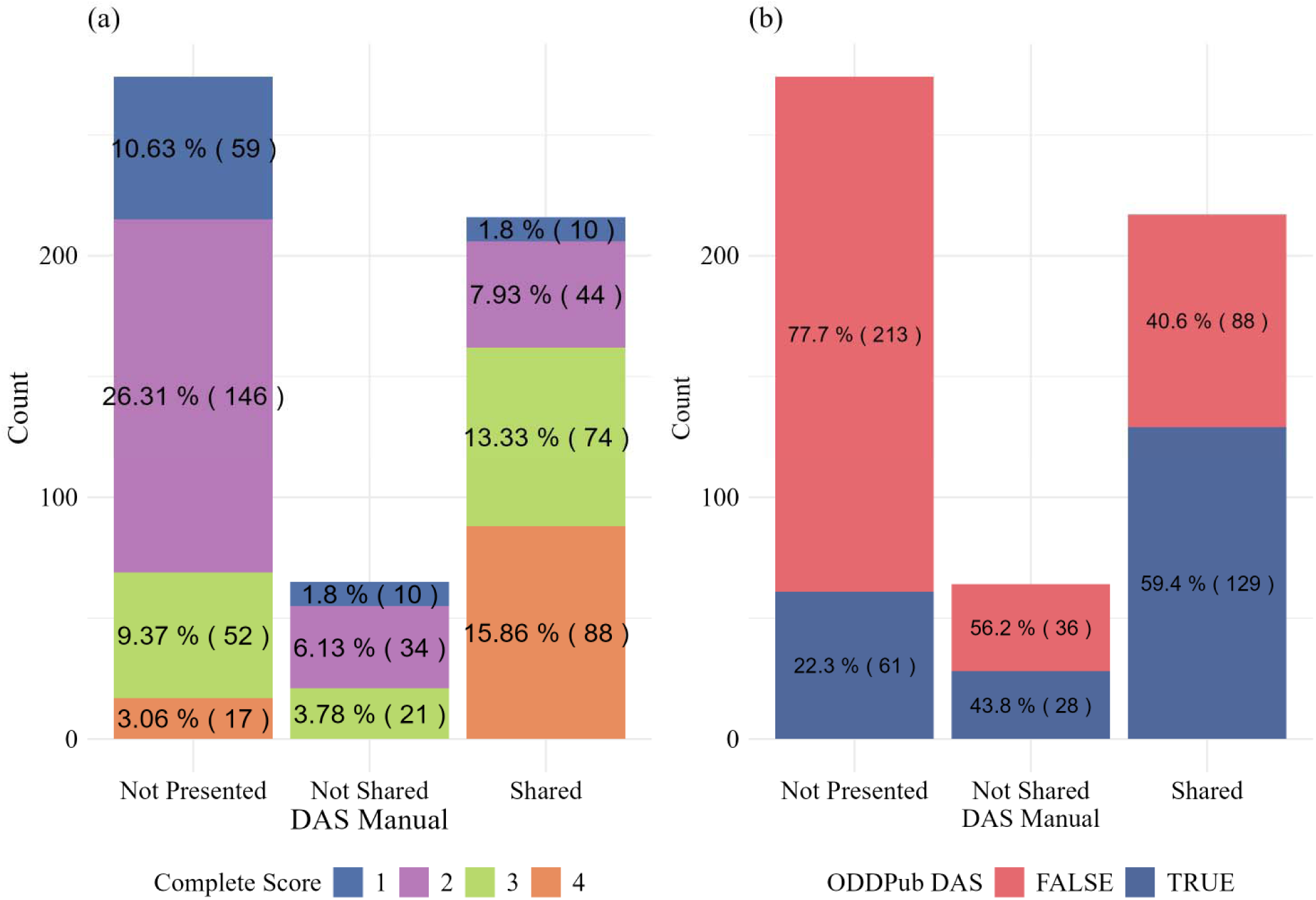
Manual and automated evaluation of Data Availability Statement (DAS) status. (a) Distribution of manually-evaluated DAS status across Completeness score categories for the 555 research publications. Most articles that shared all data (88 with Completeness 4) included a DAS signalling this Open sharing. (b) Comparison of Data Availability Statement (DAS) evaluation between the Manual assessment and ODDPub detection (extracted from open_data_category as True/False, see Supplementary Material File 2).

Further analysis reveals that the presence of a DAS, irrespective of its content, considerably enhances the detection capabilities of ODDPub. When a DAS is present, our analysis showed an increase in the sensitivity of ODDPub to 61.8% from the initial 52%, indicating an improved ability to correctly identify instances where data is indeed openly available. However, specificity saw a slight decrease to 85.71% from 90%. This change underscores that while the presence of a DAS generally improves the accuracy of detecting true positives, there may be a minor trade-off in the specificity due to the varied nature of DAS content (Supplementary Material File 2).

## Discussion

Our analysis of 555 bioscience research papers from the University of Edinburgh has shown a progressive shift towards sharing more research data over the last ten years and sharing more FAIRly. Our sample size is larger than most previous studies that assessed research data sharing within medical and health specialities, as discussed in Hamilton et al. (24). However, our colleagues’ research ranges from the atomic scale of molecular structures to populations of intact organisms, so there is no common data unit here, like ecological research (14) and in contrast to the individual participant (or patient) data of many human-centric studies (24).

Our 4-level scoring system evaluates data based on Completeness, Accessibility, Reusability, and License, accommodating diverse data types and volumes. The protocol is detailed in a repository for future reference (see Data Availability), this system may benefit from increased precision for targeted sample assessments involving data volumes. While ODDPub, as an automated text mining algorithm, effectively identifies unshared data with high specificity, its ability to accurately detect shared data is comparatively weaker. Apart from Completeness, our criteria evaluated the curation and FAIRness of each article’s best-shared dataset, with potential uniform application in future research as demonstrated using a smaller dataset (25).

Survey data suggests that research data sharing was already increasing globally by 2014 (26), when our sample starts. Earlier studies like Roche et al. (14) indicate that 56% of ecology data in the Dryad repository from 2012-2013 was incomplete (Completeness score <3). Although our data starts in 2014, we found similar trends of incompleteness, with 80% of our first year’s data scoring below 3 in completeness. In contrast, a 2019 study on global cancer research reported that only 19% of the publications made some or all their data publicly available (21). However, our 2019 sample from diverse domains, including non-communicable diseases like cancer, showed that 52% of papers shared their data (completeness >2). These variations underscore that the initial point of assessment—whether it is a data repository or publication—and the research domain can significantly influence the observed rates of data sharing.

Several policy changes have significantly influenced data sharing practices. The articulation of the FAIR principles in 2016 (6), followed by their progressive adoption, set new standards for data management. Similarly, the University of Edinburgh updated its ‘Research Data Management’ policy in 2021 (27), improving upon its 2011 guidelines to strongly advocate for research data sharing in alignment with the FAIR principles. Additionally, data-sharing mandates during the COVID-19 pandemic, which impacted the UK from 2020 (28,29), further accelerated these trends. Our results reflect these influences, showing a rapid increase in data sharing over the last three years. The proportion of articles fully sharing data nearly doubled, rising from 22% in 2021 to 45% in 2023. The highest scores for each of our criteria were achieved between 2021 and 2023.

Our manual scoring revealed significant variations in data sharing across different types. Out of 329 papers involving genomic data, 241 shared their data, benefiting from established databases like GenBank and GEO which promote a strong sharing culture among researchers, publishers, and funders (30–32). In contrast, only 69 out of 352 papers that used imaging data shared it; the BioImage Archive (BIA) only recently started accepting significant contributions (33). General data repositories like Figshare and Zenodo were commonly used, though they lack specialised features for microscopy images and struggle with large dataset uploads (>50GB) due to reliability issues (34). Due to these challenges, amongst other factors, sharing data in the article’s Supplementary Materials also remains a common practice (340 out of 555 articles). These technical and infrastructural challenges extend beyond mere data sharing; they also critically impact data storage during the research phase. Managing large image datasets, in particular, can become expensive and logistically demanding, as highlighted by recent findings (35). This reality underscores the need for more data management and infrastructural support to facilitate data handling and accessibility (36).

Sharing human data presents additional hurdles, including privacy (36,37), ethical considerations (38), and regulatory challenges (39). Of 159 papers involving human data, only 48 shared their data, highlighting a need for further exploration of this moderate sharing rate (40). The fact that many papers in our sample used cell lines may reduce privacy concerns compared to data directly from human subjects (21). Additionally, there might be varying motivations and skills among researchers within our organisational framework (41). Our Completeness score focuses on open data sharing, not considering conditional access that might still meet FAIR standards.

Research publishers’ Open data policies have significantly advanced data sharing, which we measure by the presence of explicit Data Availability Statements (DAS). Our approach, scoring these explicit statements, contrasts with other studies that assess data availability declarations regardless of their format or location within the article (42). Importantly, the presence of a DAS notably enhances the effectiveness of ODDPub, which, like other text detection tools, relies on explicit reporting. Standardising these statements can greatly improve documentation, making automated algorithms more accurate—especially as the trends move towards integrating automated algorithms as the initial step in detecting data sharing (43), whether with or without manual assessment. This accuracy is critical because any discrepancies in documentation can lead to missed instances of data sharing.

However, a DAS alone does not guarantee actual data sharing (44). In our sample, 10 papers had a DAS stating “all data were available,” yet they shared no open data, with the latest published in 2021. Sometimes, authors might confuse the visual display of data, like graphs, with sharing underlying numerical datasets. This continued gap in compliance reflects inadequate enforcement by editors and peer reviewers (45). The potential for addressing these issues was recognised in a 2023 UK Parliament inquiry (46), highlighting ongoing challenges in both policy development and implementation, including adequate repository support to meet the demands of modern research and policymaking.

## Standpoint

The authors worked either at the School of Biological Sciences, University of Edinburgh during the study, in the Biological Research Data Management team (BioRDM), or for EW and MMcL, in the Collaborative Approach to Meta Analysis and Review of Animal Experimental Studies (CAMARADES) research group in the College of Medicine and Veterinary Medicine. The BioRDM team grew from the need for data management in our interdisciplinary biology research projects in the Centre for Systems Biology at Edinburgh (later SynthSys, now Centre for Engineering Biology), since 2008. The CAMARADES group develops systematic reviews of preclinical neuroscience research. AJM also chaired the UKRI-BBSRC Review of Data-Intensive Bioscience (2020).

## Data and Code Availability Statement

All data utilised in this study, along with the detailed code, are available in the Zenodo repository dedicated to this paper (https://doi.org/10.5281/zenodo.14169649) and should be cited as (APA Style):

“*Deeb, H., Creasey, S., de Ugarte, D. L., Strevens, G., Usman, T., Yun Wong, H., Kutzer, M. A. M., Wilson, E., Zieli*ń*ski, T., & Millar, A. J.* (*2024*)*. A Decade of Progress: Open Data Practices in Bioscience at the University of Edinburgh (1.0) [Data set]. Zenodo.* https://doi.org/10.5281/zenodo.14169649.”

The main repository of the project with all the previous data and output can be found on Github: BioRDM/InsightsOfOpenPracticesInBiosciences [Internet]. BioRDM; 2024 [cited 2025 Jan 7].

Available from: https://github.com/BioRDM/InsightsOfOpenPracticesInBiosciences. The detailed scoring protocol can be retrieved and adapted from the protocols.io repository:

https://dx.doi.org/10.17504/protocols.io.kxygxyxmdl8j/v2. The preprint of the article can be found on BioRxiv.

## Supplementary Materials Captions

The supplementary materials for this paper are two files attached to the paper and available on the Zenodo repository:

- Supplementary Material File 1 - Figures and Tables: The additional figures and tables that support our results.
- Supplementary Material File 2 – ODDPub: This file provides the methodology, the variables and all the results of using ODDPub in our biological samples.

## Authors’ Contributions

AJM designed the study; HYW and TU adapted scoring criteria and performed primary scoring; GS, DLU and SC performed primary and secondary scoring; HD performed secondary scoring, analysed the data, drafted the manuscript, and shared the outputs; EW performed the ODDPub analysis; HD and MK designed the analysis methods; TZ and AJM edited the manuscript and supervised the work.

## Acknowledgments

We extend our sincere gratitude to Professor Simon N. Wood, Chair of Computational Statistics at the School of Mathematics, and his PhD student, Antoni Sieminski, for their invaluable support and expert consultation provided through University of Edinburgh statistics drop-in clinics. We also wish to thank Professor Malcolm MacLeod for his research supervision for EW. Additionally, we would like to acknowledge the University of Edinburgh Library Research Support team for their assistance in accessing the information systems that were crucial in identifying our colleagues’ publications.

