## Supplementary Material File 1 for "The Rise of Open Data Practices Among Bioscientists at the University of Edinburgh"

### **Supplementary Figure**

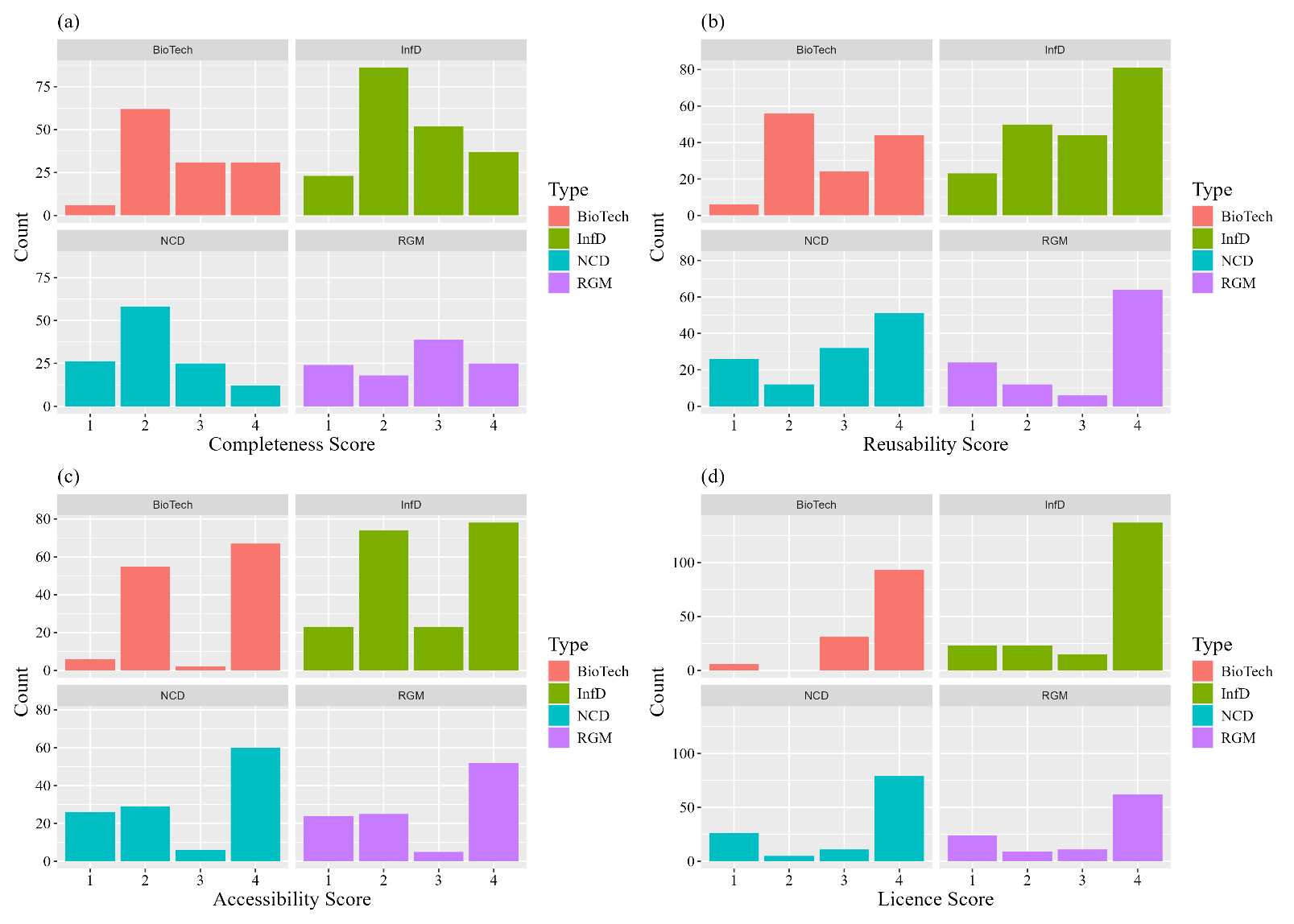

**Figure S1**: *Distribution of the four criteria among the research types.*

The distribution of Completeness (a), Reusability (b), Accessibility (c), and License (d) scores by research area. Research groups were selected in four areas: Non-Communicable Diseases (NCD, green), Infectious Diseases (InfD, blue) and their treatments, Microbial Biotechnology (Biotech, red), Stem Cells and Regenerative Medicine (RGM, purple).

### **Supplementary Tables:**

| **File Extension** | **Non-Proprietary** | **Human-readable** | **Machine-readable** |
| --- | --- | --- | --- |
| .ab1 | 1 | 0 | 1 |
| .avi | 0 | 1 | NA |
| .bam | 1 | 0 | 1 |
| .csv | 1 | 1 | 1 |
| .doc | 1 | 1 | 0 |
| .docx | 1 | 1 | 0 |
| .fastq | 1 | 1 | 1 |
| .gif | 0 | 1 | NA |
| .jpg | 1 | 1 | NA |
| .maf | 1 | 1 | 1 |
| .mov | 0 | 1 | NA |
| .mp4 | 0 | 1 | NA |
| .pdf | 0 | 1 | 0 |
| .raw* | 0 | 1 | 1 |
| .rtf | 0 | 1 | 0 |
| .sas | 0 | 1 | 1 |
| .sav | 0 | 1 | 1 |
| .tif | 1 | 1 | NA |
| .txt | 1 | 1 | 1 |
| .vcb | 1 | 1 | 1 |
| .wav | 0 | 1 | 0 |
| .xls | 1 | 1 | 1 |
| .xlsx | 1 | 1 | 1 |
| .xml | 1 | 1 | 1 |
| **Table S1**: Characteristics of data file formats.  “0” represents no, “1” is yes. NA was regarded as “yes” when scoring for “Data Reusability”.  This table was adapted from Roche et al. (14).  * .raw, A camera raw image file contains unprocessed or minimally processed data from the image sensor of either a digital camera, a motion picture film scanner, or other image scanner. | | | |

|  | **Completeness** | **Reusability** | **Accessibility** | **Licence** |
| --- | --- | --- | --- | --- |
| Year | 1.21 (1.14 - 1.28) *** | 1.21 (1.15 - 1.28)*** | 1.19 (1.13 - 1.26) *** | 1.11 (1.05 - 1.18)*** |
| Year+ RE | 1.24 (1.17-1.32) *** | 1.22 (1.15 – 1.30)*** | 1.20 ( 1.13-1.27)*** | 1.13 (1.05-1.20)*** |
| **Table S2**: Influence of Year of publication on Scoring Criteria.  Ordinal regression analysis without, or with (+RE), the random effect of research groups for each Criterion.  Notes:   - The numbers presented are Odds Ratios (OR) with 95% Confidence Intervals (CI). - Statistical significance is indicated by: *p-value < 0.05, **p-value < 0.01, and ***p-value < 0.001. | | | | |

|  | **Biotechnology** | **Infectious Diseases** | **Non-Communicable diseases** | **Regenerative Medicine** |
| --- | --- | --- | --- | --- |
| Completeness | Reference | 0.75 (0.50 – 1.11) | 0.40 (0.26 – 0.64)*** | 0.95 (0.59-1.52) |
| Completeness +RE | Reference | 0.72 (0.41 – 1.26) | 0.37 (0.20 – 0.70)*** | 0.88 (0.46-1.70) |
| Reusability | Reference | 1.21 (0.82-1.78) | 1.19 (0.76 – 1.85) | 1.78 (1.09 – 2.93)** |
| Reusability +RE | Reference | 1.20 (0.78-1.83) | 1.18 (0.73 – 1.91) | 1.76 (1.03 – 3.00)** |
| Accessibility | Reference | 0.70 (0.46 – 1.05) | 0.75 (0.47-1.19) | 0.72 (0.44 – 1.17) |
| Accessibility +RE | Reference | 0.68 (0.39 – 1.18) | 0.74 (0.39-1.38) | 0.64 (0.33 – 1.22) |
| Licence | Reference | 0.75 (0.47 – 1.21) | 0.59 (0.35 – 0.99)** | 0.45 (0.26 – 0.76)*** |
| Licence +RE | Reference | 0.75 (0.40 – 1.38) | 0.57 (0.29 – 1.12) | 0.41 (0.20 – 0.82)** |
| **Table S3**: Influence of research area on Scoring Criteria.  Ordinal regression analysis without, or with (+RE), the random effect of research groups for each Criterion.  Notes:   - The numbers presented are Odds Ratios (OR) with 95% Confidence Intervals (CI). - Statistical significance is indicated by: *p-value < 0.05, **p-value < 0.01, and ***p-value < 0.001. | | | | |

| **Variable** | | **Bio-technology** | **Infectious Diseases** | **Non-Communicable diseases** | **Regenerative Medicine** | **Total** |
| --- | --- | --- | --- | --- | --- | --- |
|  |  | (n= 130) | (n=198) | (n=121) | (n=106) | (n=555) |
| Genomic Data | Used and Shared | 27 (20.77%) | 72 (36.26%) | 42 (34.71%) | 100 (94.34%) | 241 (43.42%) |
|  | Used and not Shared | 27 (20.77%) | 25 (12.63%) | 30 (24.79%) | 6 (5.66%) | 88 (15.86%) |
|  | Not Used | 76 (58.46%) | 101 (51.01%) | 49 (40.50%) | 0 (0.00%) | 226 (40.72%) |
| Image shared | Used and Shared | 11 (8.46%) | 19 (9.60%) | 8 (6.61%) | 31 (29.24%) | 69 (12.43%) |
|  | Used and not Shared | 57 (43.84%) | 79 (39.90%) | 84 (69.42%) | 63 (59.43%) | 283 (51.00%) |
|  | Not Used | 62 (47.69%) | 100 (50.0%) | 29 (23.97%) | 12 (11.32%) | 203 (36.58%) |
| Human Data | Used and Shared | 0 (0.00%) | 27 (13.64%) | 14 (11.57%) | 7 (6.60%) | 48 (8.65%) |
|  | Used and not Shared | 0 (0.00%) | 27 (13.64%) | 43 (35.54%) | 41 (38.68%) | 111 (20.00%) |
|  | Not Used | 130 (100%) | 144 (72.73%) | 64 (52.89%) | 58 (54.72%) | 396 (71.35%) |
| Data Storage | Repository | 18 (13.85%) | 47 (23.74%) | 48 (39.67%) | 11 (10.37%) | 124 (22.34%) |
|  | Supplementary | 69 (53.08%) | 75 (37.88%) | 42 (34.71%) | 21 (19.81%) | 207 (37.30%) |
|  | Both | 37 (28.46%) | 45 (22.73%) | 3 (2.48%) | 48 (45.28%) | 133 (23.96%) |
|  | DOIs in Methods | 0 (0.00%) | 8 (4.04%) | 2 (1.65%) | 2 (1.89%) | 12 (2.16%) |
|  | No Storage | 6 (4.62%) | 23 (11.62%) | 26 (21.49%) | 24 (22.64%) | 79 (14.23%) |
| **Table S4**: Data Sharing Practices in four Research Areas. | | | | | | |

| **Sharing Variables** | | **Completeness** | **Reusability** | **Accessibility** | **Licence** |
| --- | --- | --- | --- | --- | --- |
| Data Availability Statement (DAS) | DAS with Open Data | 9.28  (6.08 - 14.16)*** | 7.98  (5.34 - 11.94)*** | 8.62  (5.65 - 13.17)*** | 6.06  (3.66 - 10.02)*** |
|  | DAS with Data upon request | 1.46 (0.86 – 2.46) | 2.60  (1.57 - 4.32)*** | 2.43 (1.42 – 4.14)*** | 2.24  (1.22 – 4.11)*** |
|  | Not Presented | Reference Group | | | |
| Preprint | Yes | 1.86 (1.22-2.83)** | 1.33 (0.87-2.04) | 1.51 (0.96 – 2.39) | 1.40 (0.81 - 2.42) |
|  | No | Reference Group | | | |
| **Table S5**: Influence of Data Availability Statement (DAS) and Preprint Status on Scoring Criteria adjusted with the random effect of research groups for each Criterion.  Ordinal regression is used to analyse the scores relative to the reference group of articles without a DAS, or without a preprint, including the potential influence of the Research Group as a random effect.  OR interpretation: OR values reflect the multiplicative increase in the odds of achieving higher criterion scores, where each unit of DAS categories or the presence of a preprint can elevate the odds by up to several times, depending on OR of the specific criterion assessed.  Note:   - The numbers presented are Odds Ratios (OR) with 95% Confidence Intervals (CI). - Statistical significance is indicated by asterisks:   *p-value < 0.05, **p-value < 0.01, and ***p-value < 0.001. | | | | | |
