## Supplementary Material File 2 for "The Rise of Open Data Practices Among Bioscientists at the University of Edinburgh"

**ODDPub Supplementary Material File**

**Introduction**

Significant challenges continue to impede the systematic monitoring and quantification of adherence to open data practices within biomedical literature. This complication primarily arises from the variability in reporting standards and the overwhelming volume of publications, factors that render manual assessments both labor-intensive and susceptible to inconsistency.

Acknowledging the urgent need for scalable and dependable tools to evaluate open data practices, Riedel et al. (2020) (1) have innovatively developed Open Data Detection in Publications (ODDPub), a sophisticated dictionary-based text mining algorithm tailored to automate the detection of open data sharing occurrences in biomedical publications. The introduction of ODDPub represents a critical progression in the field, empowering researchers and policymakers with a high-throughput approach to systematically screen large volumes of publications for statements of open data and references to shared research data.

The decision to incorporate ODDPub into our analysis framework originates from the desire to corroborate and expand upon our initial manual evaluations of data-sharing practices observed in our dataset. While these manual assessments provide detailed insights, they are inevitably limited by the enormous dataset scope, the potential for subjective discrepancies and the practical bounds of manual paper analysis capabilities.

The ODDPub algorithm is designed to be used in R programming software. We retrieved publication full texts in PDF format from DOIs using the Find Full Text function in EndNote 20 (RRID:SCR_014001) or handsearching and imported these into an R Project environment. We used the pdftools R package (2) to convert PDF files into text format for processing by ODDPub. ODDPub extracted the sentence relevant to data / code sharing and assigned each publication a binary (TRUE/FALSE) score each for the presence of data sharing and the presence of code sharing. Additionally, ODDPub categorised the type of data sharing (e.g. supplement, general-purpose repository, field-specific repository).

**Assessing ODDPub Outcomes Through Sensitivity and Specificity Metrics**

In the context of evaluating the performance of the ODDPub algorithm using sensitivity and specificity, the terms True Positives (TP), False Negatives (FN), True Negatives (TN), and False Positives (FP), in relation to both the algorithm's detection results and the manual assessments, were used.

- True Positives (TP):

Definition: These are cases where ODDPub correctly identifies open data/code as present, and this detection aligns with the findings from manual assessments (i.e., the manual assessment also confirms that open data/code is present).

- False Negatives (FN):

Definition: These occur when ODDPub fails to identify the presence of open data/code, but the manual assessment indicates that open data/code is indeed present.

- True Negatives (TN):

Definition: These are cases where ODDPub correctly identifies that there is no open data/code, and the manual assessment confirms that open data/code is indeed not present.

- False Positives (FP):

Definition: These occur when ODDPub identifies open data/code as present when it is not, according to the manual assessment.

**Calculation of Sensitivity and Specificity:**

- **Sensitivity (True Positive Rate):**

Sensitivity=True Positives (TP) / (True Positives (TP) + False Negatives (FN)

Interpretation: Sensitivity measures the proportion of actual positive cases of open data/code presence that are correctly identified by ODDPub compared to the manual assessment. It reflects the algorithm’s ability to detect open data/code when it indeed exists.

- **Specificity (True Negative Rate):**

Specificity = True Negatives (TN) / True Negatives (TN) + False Positives (FP)

Interpretation: Specificity measures the proportion of actual negative cases (where there is no open data/code) that are correctly identified by ODDPub in alignment with the manual assessment. It indicates the algorithm’s ability not to falsely label non-existing open data/code as present.

**Section One: Open Data Analysis**

ODDPub identifies open data using a dictionary-based text mining approach and scores publications with a binary TRUE/FALSE evaluation system, where data qualifies as open if it is shared alongside the respective publication and meets specific conditions. These conditions include the data being either raw or preprocessed to enable analytical replication (echoing our criterion of completeness with a required score of 3 or 4), originating from primary or assembled secondary sources authored by the publication's authors, and formatted in a machine-readable manner (aligning with our reusability criterion with scores of 3 or 4). Additionally, ODDPub mandates that data must be easily accessible without restrictions such as registration or special access (paralleling our accessibility criterion with scores of 3 or 4), and ideally deposited in recognised repositories to enhance findability and reuse potential.

In contrast, our criteria are segmented into four detailed dimensions: completeness, reusability, accessibility, and licensing, offering a more nuanced evaluation framework compared to ODDPub's approach. To ascertain alignment with ODDPub’s definition, we rate datasets on these dimensions with a variable called “Manual Open Data”. A dataset is classified as "TRUE" for open data if it achieves a completeness score of 3 or 4; it can also earn a "TRUE" if the completeness is slightly lower (score of 2), but reusability or accessibility scores are at 3 or 4, reflecting a compensatory balance among the criteria. Conversely, datasets that do not meet these threshold combinations are designated as "False." This methodology ensures a rigorous yet flexible analysis, harmonising our intricate criteria with ODDPub's more streamlined, binary classification to robustly assess the open data landscape in our studies.

a- **ODDPub vs Completeness only**

| **Table O1 (a)** |  | **ODDPub Open Data** | |  |
| --- | --- | --- | --- | --- |
| **Completeness** |  | TRUE | FALSE | Total |
|  | 1 | 8 | 71 | 79 |
|  | 2 | 57 | 167 | 224 |
|  | 3 | 80 | 64 | 144 |
|  | 4 | 73 | 35 | 108 |
|  | total | 218 | 337 | 555 |

| **Table O1 (b)** |  | **ODDPub Open Data** | |
| --- | --- | --- | --- |
|  |  | TRUE | FALSE |
| **Completeness=1** | FALSE | 8 | 71 |
| **Completeness>1** | TRUE | 210 | 266 |

***Table O1 a & O1 b:*** *Distribution of TRUEand FALSEOpen Data Detections from ODDPub Across Different Scores of Completeness.*
**Green cells indicate instances where the program correctly matches the manual assessment.*
**Red cells signify instances where the program did not match the manual assessment.*

In comparing ODDPub's open data determination with our completeness score, the sensitivity is observed at 44.12%, indicating it correctly identifies about 44% of true positives. The specificity stands at 89.87%, showing high accuracy in avoiding false positives.

b- **ODDPub vs. The Four Criteria of the Manual Assessment (Completeness, Resuability, Accessibility, and Licence)**

|  |  | **ODDPub Open Data** | |  |
| --- | --- | --- | --- | --- |
|  | **Row Labels** | **FALSE** | **TRUE** | **Total** |
| Manual Open Data | False | 147 | 17 | 164 |
|  | True | 190 | 201 | 391 |
|  | **Grand Total** | **337** | **218** | **555** |

*Table O2: Distribution of True and False Open Data Detections from ODDPub versus Manual Open Data scoring, incorporating all the criteria*

In this context, the Sensitivity is 52% and the specificity 90%. The improvement in sensitivity (52%), when compared to the completeness-only assessment (44.12%), indicates that integrating additional criteria (Reusability, Accessibility, Licensing) alongside Completeness improves the strength of the ODDPub evaluation. This refined approach not only supports a greater detection of true open data instances but also maintains a high level of specificity (90%).

The ODDPub paper initially presented a sensitivity of 73% and a specificity of 97% in its validation set, underscoring a highly precise method for identifying true negatives while still capturing a significant majority of true positives. In our evaluation, ODDPub demonstrated a specificity of 89.87%, closely aligning with the original paper's findings, confirming its robustness in accurately rejecting non-open data instances. However, our observed sensitivity of 44.12% for ODDPub is notably lower than that reported in the original study, suggesting a possible variance in the datasets or differences in the operational definitions of open data used in our analysis compared to the original ODDPub settings.

**Section Two: Open Code Analysis**

In this section, we discuss the evaluation of open code-sharing practices, examining the alignments and discrepancies between our manually categorised data and the automated ODDPub detection. Our manual categorisation differentiates between 'Code Not Shared', 'Code Shared', and situations where 'No Code Needed' to produce the research results. This classification underscores a detailed approach to coding availability, recognising cases where coding is irrelevant to the research outcomes—a distinction not made by the ODDPub algorithm, which operates on a binary TRUE/FALSE detection system.

| **Table O3 (a) ODDPub Code** | | | | |
| --- | --- | --- | --- | --- |
|  |  | **FALSE** | **TRUE** | **Total** |
| **Code Variable -  Manual** | Code Not Shared (False) | 75 | 4 | 79 |
|  | Code Shared (True) | 79 | 33 | 112 |
|  | No Code Needed | 356 | 8 | 364 |
|  | **Total** | **510** | **45** | **555** |

*Table O3a: Comparison of Manual Coding Classification and ODDPub Detection*

| **Table O3 (a)** |  | **ODDPub Code** | |
| --- | --- | --- | --- |
|  |  | **FALSE** | **TRUE** |
| **Code Variable - Manual** | **TRUE** | 79 | 33 |
|  | **FALSE** | 431 | 12 |
|  | **Total** | 510 | 45 |

*Table O3b: ODDPub Code Detection Compared to Manual Truth*

From the data illustrated in Table 3b, the sensitivity and specificity of the ODDPub algorithm in detecting open code are quantitatively assessed. The sensitivity, at 29.46%, indicates that ODDPub correctly identifies just 30% of the true positives, suggesting lower effectiveness in accurately pinpointing cases where code is shared. However, the specificity is exceptionally high at 97.29%, demonstrating ODDPub’s robust capability in correctly identifying negatives—cases where the code is not shared or not needed.

**Section Three: Data Availability Analysis.**

In this section, we focus on how Data Availability Statements (DAS) are evaluated within our study compared to their treatment in the ODDPub framework. Our evaluation categorises DAS into three distinct types:

- DAS with Restrictive: Indicates the presence of a DAS, but the data is not openly shared.
- DAS Open: Implies that a DAS is present and the data are openly available and shared.
- No DAS: Signifies that no specific DAS is mentioned regardless of the data-sharing status.

Unlike our structured approach, ODDPub does not directly categorise data availability statements into specific categories related to restrictions or openness. Instead, it uses the open_data_category variable to distinguish whether data is shared through specific or general repositories or supplementary materials, which gives an indirect indication of the presence of some form of DAS.

To align ODDPub’s methodology with our criteria for evaluating DAS, we constructed a new variable termed ODD_DAS. This variable is assigned a value of "Yes" if the open_data_category from ODDPub indicates a clear DAS is mentioned (regardless of the data sharing specifics like the type of repository). Conversely, "No" is assigned if the open_data_category lacks any explicit mention of a DAS, even if the category suggests the data might be in a repository. This distinction is crucial as it focuses on the presence of a statement about data availability, rather than the location or openness of the data itself, aligning more closely with how DAS is assessed in our methodology.

|  |  | **ODDPub_DAS** | |  |
| --- | --- | --- | --- | --- |
|  |  | **No** | **Yes** | **Total** |
| **Manual DAS assessment** | DAS Restrictive | 52 | 12 | 64 |
|  | DAS Open | 160 | 57 | 217 |
|  | Not presented | 268 | 6 | 274 |
|  | **Total** | **480** | **75** | **555** |

*Table O4a: Comparison of Manual DAS Classification and ODDPub Detection*

|  |  | **ODDPub_DAS** | |
| --- | --- | --- | --- |
|  |  | TRUE | FALSE |
| **Manual DAS assessment** | TRUE | 69 | 212 |
|  | FALSE | 6 | 268 |

Table O4b: ODDPub DAS Detection Compared to Manual Truth

The sensitivity of the ODDPub_DAS is approximately 24.56%, demonstrating that it identifies about a quarter of true positive cases where a DAS is indeed present. Despite the low sensitivity, the specificity reached a high value of 97.81%, indicating that the program effectively recognises the absence of a DAS, with a very low rate of false positives. In practical terms, this means that we can trust the program at a 98% confidence level to correctly detect papers where a DAS does not exist.

A key challenge identified in the ODDPub's DAS detection capability is its restrictive focus on reading explicit mentions labeled precisely as "Data Availability Statement." The heterogeneous nature of data-sharing declarations in academic literature often encompasses a variety of titles, such as "Data and Code Availability," "Methods and Data Availability," "Open Research Statement," or "Open Data Statement." This variation in terminology can lead to substantial under-detection of valid DAS, contributing to the observed low sensitivity of the program.

As we conclude our analysis of DAS through both manual assessment and automated detection via the ODDPub system, it is evident that while ODDPub excels in confirming the absence of DAS, it is less effective at identifying the presence of a varied array of DAS titles and forms. This highlights a potential area for future refinement, suggesting that extending the algorithm to recognise a broader spectrum of DAS-related terminology could significantly enhance its practical utility and sensitivity.

**Section 4: Effect of Data Availability Statements on the Accuracy of Open Data Detection**

This section examines how the presence of Data Availability Statements (DAS), regardless of whether they are restrictive or facilitate open data, affects the sensitivity and specificity of ODDPub in accurately detecting open data access. To accomplish this, we filtered our original dataset to include only those studies that featured any form of DAS, analysing how standardisation in reporting such as the mere presence of a DAS influences detection performance.

|  |  | **ODDPub Open Data** | |  |
| --- | --- | --- | --- | --- |
|  |  | **FALSE** | **TRUE** | **Total** |
| **Open Data Manual** | F | 30 | 5 | 35 |
|  | T | 94 | 152 | 246 |
|  | **Total** | **124** | **157** | **281** |

Table O5: Impact of Data Availability Statement Presence on Open Data Detection

Post-filtering, the sensitivity and specificity of detecting open data were recalculated:

- Sensitivity: Increased to 61.79% from the original 52%, indicating an improved ability to identify true positives (i.e., correctly identifying cases where data was indeed open).
- Specificity: Reduced slightly to 85.71% from the original 90%, still reflecting a capacity to identify true negatives correctly, even though a minor decrement.

The observed improvements in sensitivity highlight the beneficial impact of standardised data reporting practices, such as DAS. The presence of a DAS, regardless of its content, significantly enhances the model’s precision in identifying open data cases. This suggests that standardisation, even at a basic level of reporting presence, can aid significantly in improving the detection algorithms employed by ODDPub.

**Conclusion:**

Our analysis of the ODDPub algorithm's performance in detecting open data and code sharing practices reveals a notable disparity between its specificity and sensitivity. The results clearly show that the algorithm excels in identifying the absence of data or code sharing, demonstrating high specificity. However, it exhibits lower sensitivity in recognising instances where data or code is actually shared. This discrepancy likely stems from the varied ways researchers report their sharing practices, which can challenge text-based detection algorithms that rely on specific cues to identify openness.

Given these findings, it is imperative to advocate for standardised reporting practices within the scientific community. Uniformity in how data and code sharing are documented will not only enhance the accuracy of current algorithms but will also facilitate the development of more sophisticated systems capable of reliably recognising open practices. As the landscape of open science evolves, fostering clarity and consistency in reporting will undoubtedly benefit the efficacy of automated tools designed to promote transparency and reproducibility in research.
